# Matrix effects influence biochemical signatures and metabolite quantification in dried blood spots

**DOI:** 10.1101/2025.07.22.666056

**Authors:** Jan-Clemens Cremer, Viktorija Juric, Jakob Koch, Sabine Scholl-Bürgi, Daniela Karall, Johannes Zschocke, Markus A. Keller

## Abstract

Dried blood spots (DBS) represent a convenient clinical sample material, offering low infection risk, easy transport, and long-term metabolite stability. However, applying samples such as whole blood, serum or plasma onto filter paper introduces an additional matrix, potentially affecting metabolite extraction. Here, we compare metabolite recovery from liquid samples and their filter paper analogue using both targeted (acylcarnitines, amino acids) and untargeted metabolomics. Significant matrix effects were observed for some compounds, especially for dicarboxylic acylcarnitines (C3DC–C6DC) and specific amino acids (cystine, cystathionine). We did not identify specific metabolite characteristics that may predicted altered recovery. In a cohort of 229 authentic DBS samples — including patients diagnosed with inherited metabolic disorders, obesity or under a ketogenic diet — untargeted profiling combined with random-forest machine learning led to an effective stratification. Notably, C4DC, despite strong matrix effects, was ranked in the top ten variables of this random-forest model. With adequate validation, DBS can be safely used for diagnostic purposes despite possible matrix effects, but care must be taken in the comparison of values obtained when different sample materials are used.

## Introduction

Dried blood spots (DBS) on standard filter paper cards are a convenient sample type in a diverse spectrum of clinical applications (1, 2). Their utility became apparent in the early 1960’s when they were used by Guthrie et. al. in newborn screeing for phenylketonuria. The original test was a bacterial growth inhibition assay that allows identification of elevated phenylalanine levels (3). Liquid chromatography coupled with tandem mass spectrometry (LC-MS/MS) has become the preferred analytical technique for this and other applications because of superior flexibility, through-put and quantifiability, as well as reduced costs (4), but DBS has remained the sample matrix of choice. Additionally, DBS have found applications in connection with other meaningful readouts such as ELISA and immunoblotting (5–7). DBS hold several advantages over other biological sample types such as serum, plasma or whole blood, including a low risk of infection, easy sample handling, transport and storage (8). However, disadvantages of DBS include error-prone assessment of sample volume to pursue accurate quantifications. Furthermore, the DBS characteristics are complex due to factors like haematocrit effect, analyte concentration differences between inner and outer circle punches, blood spreading, paper saturation and the impact of temperature and humidity on analyte stability (9, 10).

Today, DBS are still best known for their use in newborn screening for the early detection of endocrine and inherited metabolic disorders (IMDs) (11–14). IMDs result from genetic perturbations that alter metabolic pathways and change the composition of specific parts of the metabolome. Most IMDs are inherited in an autosomal recessive manner, although autosomal dominant and X-linked forms also occur. The overall incidence is estimated at approximately 1:1000 live births, although this number varies to a certain extent from region to region (15, 16). Various classification systems emerged over time to structure the large number of IMDs, using the clinical manifestations or pathomechanisms as categories (17). A novel approach shows a hierarchical system to group almost 1500 IMDs according to the diverse metabolic pathways that are primarily affected (18). Sampling for the diagnosis of IMDs thus must account for a wide range of metabolite types and biochemical characteristics.

Using targeted mass spectrometry it has been shown that polar compounds like acylcarnitines and amino acids together with succinylacetone can be analyzed from DBS in a single run using hydrophobic interaction liquid chromatography (HILIC), even with small punch sizes of only 3.2 mm (19, 20). In recent years, with the advancement of technological options, a strong trend towards untargeted metabolomics has emerged, where multiple chemical classes of analytes are detected in a single run (21). Untargeted metabolomics aims for a more comprehensive molecular understanding of disease phenotypes based on the potential identification of novel biomarkers or secondary dysregulated metabolites (22–24). Patient-derived DBS thus represent a valuable tool to obtain a deeper understanding for the dysregulation of metabolomic profiles in IMD (25).

One distinctive drawback of untargeted metabolomics is the near impossibility to incorporate internal standards for all detectable compound classes in the study design (26). Thus, analytical results can be highly susceptible to matrix effects, depending on the specific compound class studied (27). Upon application to the filter paper the sample interacts with chemically diverse constituents, which facilitates potential interactions and binding events. Furthermore, the drying phase of the DBS and its exposure to atmospheric oxygen may accelerate or enable chemical transformations within the sample (28). As a result, a range of metabolic changes may occur that do not represent the sample content in the individual case but arise from the sample matrix itself.

In this study, we evaluate potential limitations that arise from the interaction between filter paper and liquid biological samples. Using targeted and untargeted metabolomics, we separately assess the filter paper-dependent matrix effects for serum, plasma, and whole blood samples. Furthermore, we assess applicability of untargeted metabolomics through the analysis in a DBS cohort of more than 200 IMD patients and controls.

## Materials and Methods

### Reagents

13 deuterated acylcarnitines (NSK-B-1, Eurisotop, Saint-Aubin, France), three further deuterated acylcarnitines (Merck, Darmstadt, Germany) and 20 isotope labelled amino acids (MSK-CAA-1, Eurisotop, Saint-Aubin, France) were used as internal standards. Acetonitrile (ACN), methanol, isopropanol (all HPLC grade) and H2O (LC-MS grade) were sourced from Merck (Darmstadt, Germany). Ammoniumformiate (NH4COOH) and formic acid (FA) were from VWR (Radnor, PA, USA). 903 DBS cards were used as filter paper (Eastern Business Forms, Greenville, SC, USA).

### Instruments

The LC-MS system consisted of either an Elute UHPLC (Bruker Daltonics, Bremen, Germany) or a Vanquish Flex UHPLC (Thermo Fisher Scientific, Waltham, MA, USA) which was coupled to a timsTOF Pro mass spectrometer (Bruker Daltonics, Bremen, Germany). Experiments were conducted with an ESI- or VIP-HESI-source in positive ionization mode and with TIMS disabled. Mass calibration was performed using a 5 mM sodiumformiate solution, which was also co-injected in each sample to enable recalibration.

### Chromatographic conditions

For targeted analysis of acylcarnitines, analytes were separated on an Acquity UPLC BEH HILIC column (2,1 x 100 mm, 1.7 µm particle size, Waters, Milford, MA, USA), while for amino acids an Acquity UPLC BEH Amide column (2,1 x 100 mm, 1.7 µm particle size, Waters, Milford, MA, USA) was used. Columns were equipped with their respective VanGuard pre-columns (2,1 x 5 mm, 1.7 µm particle size). Mobile phase A consisted of H_2_O, while mobile phase B was ACN:H2O (90:10 v/v) (both containing 0.1% FA and 5 mM NH4COOH). Details on chromatographic gradients are given in Supplementary Tables S1 and S2.

### Mass spectrometric analysis

Targeted analysis of acylcarnitines utilized the ESI-source (4,5 kV, 250°C dry gas temperature), operated within a scan range of 50 – 500 m/z. MS2 fragment spectra were obtained in bbCID mode (50.0 eV). For targeted amino acid analysis, the ESI-source was operated in the same manner, however the scan range was adjusted to 50 – 350 m/z together with slightly modified tuning settings. Detailed MS parameters are given within the supplementary material (Table S3).

Untargeted analysis of metabolites was performed either with a standard metabolomics method (ESI-source) or with optimized parameters (VIP-HESI-source). Both methods were operated within a scan range of 20 – 1,300 m/z and a capillary voltage of 4,5 kV. MS2 spectra were obtained utilizing Auto MS/MS mode (cycle time = 0.5 s). Complete method details, including tuning parameters, are given within the supplementary material (Table S3 for the standard, Table S4 for the optimized method).

### Method validation

The targeted acylcarnitine and amino acid methods were both analytically validated by assessing accuracy, precision, linearity, analytical selectivity, robustness, limit of detection (LOD), limit of quantification (LOQ), carry-over and recovery. Isotopic labelled internal standards (ISTD) were used for absolute quantification of analytes. The validation was performed with serum quality control (QC) samples purchased from ERNDIM (Manchester, UK). Validation details are listed within the supplementary material (Text S5 for acylcarnitines, Text S6 for amino acids).

### Sample preparation and experimental setup

The experimental setup for determining metabolite recovery effects from filter paper included the two extraction routes A and B (Fig. 1A). For route A, a liquid sample volume of 11 µL was extracted. For route B, the same amount of liquid sample was placed onto empty pre-cut 903 filter paper spots (6.00 mm diameter) and subsequently dried for five hours. Each analysis was performed with five technical replicates (n = 5) and the amount of total extraction solvent was adjusted to account for the liquid phase already present in route A. Serum samples (ERNDIM, Manchester, UK) for acylcarnitines and amino acids were available in two concentration levels (Level 1 = normal, Level 2 = high) and only metabolites specified on the certificate of analysis were analyzed. Authentic lithium-heparin (li.-hep.) whole blood and li.-hep. plasma were also included in the analysis. All samples were stored at -80°C until analysis. Compound nomenclature was according to supplementary Table S7. The untargeted metabolomics comparison (data processing given in supplementary Text S8) between liquid and filter paper extracts was performed with plasma and whole blood. Extracts from these samples, already prepared with the acylcarnitine method (described below), were simply remeasured using the standard metabolomics method.

**Figure 1:**
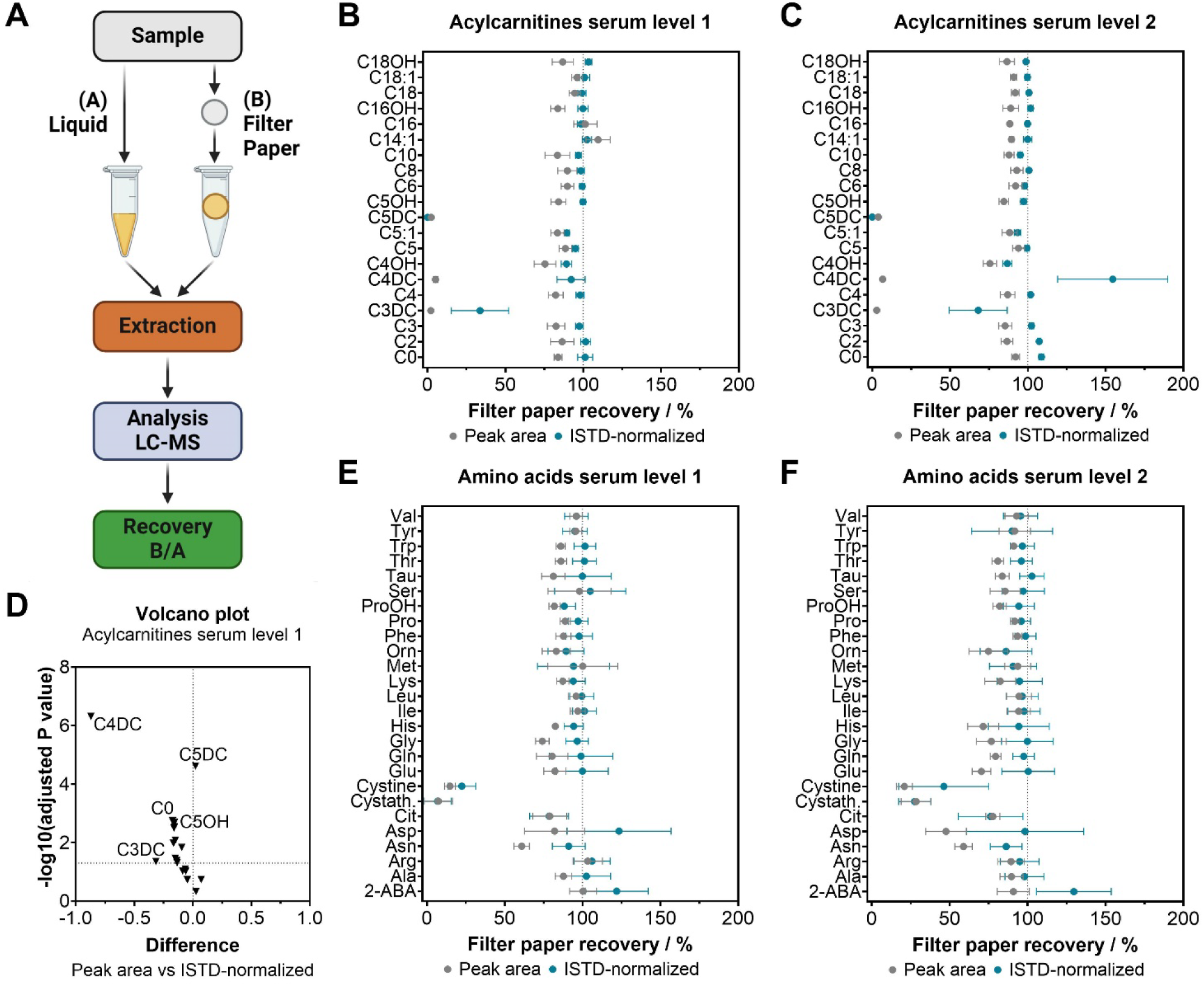
Recovery of acylcarnitines and amino acids from filter paper with artificial spiked serum. (A) Experimental setup. (B) Vertical superimposed point plots depict the mean ± SD peak area recovery and ISTD-normalized recovery of acylcarnitines in level 1 (normal metabolite concentration) serum. (C) Same as B, but for level 2 (high metabolite concentration) serum. (D) A volcano plot highlights significant differences (by multiple t-tests with Holm-Sidak correction) of peak area vs ISTD-normalized recovery for compounds from acylcarnitine level 1 serum. (E) Mean ± SD results for amino acids recovery in level 1 serum. (F) Same as E, but for level 2 serum. Recovery values are detailed in the supplementary material (File 1).

Acylcarnitines were extracted using [ACN:MeOH] (3:1 v/v) and following a protocol of shaking (20 Hz, 2.5 min), ultrasonification (10 min), shaking (30 min), centrifugation (31150 x g, 4°C, 10 min), transfer of the supernatant to autosampler vial, evaporation (under N2-flow) and subsequent resuspension (100 µL extraction solvent). From this solution, 10 µL were injected into the LC-MS system using the acylcarnitine method.

For the analysis of amino acids, [MeOH] was used as extraction solvent following a protocol of shaking (20 Hz, 2.5 min), ultrasonification (5 min), centrifugation (31150 x g, 4°C, 3 min) and transfer of supernatant to autosampler vial. From this extract, 5 µL were injected into the LC-MS system using the respective amino acid method.

### Samples

Li.-hep. whole blood was sourced from a healthy male donor (with written consent), and li.- hep. plasma was obtained accordingly by centrifugation (2000 x g, 15 min).

The clinical cohort for untargeted metabolomics analysis consisted of 229 DBS samples, which included 199 patient samples and 30 healthy controls. Clinical indications were obesity, ketogenic diet, phenylketonuria (PKU), propionic aciduria (PA), medium-chain/long-chain/very-long-chain acyl-CoA dehydrogenase deficiency (MCADD/LCHADD/VLCADD), and the carnitine-palmitoyltransferase 1 (CPT1) and 2 (CPT2) deficiencies. Due to the low number of cases, CPT1 and CPT2 were grouped together (Table 1). For the correct measurement setup, the *guidelines and considerations for untargeted clinical metabolomics studies* by Broadhurst et. al. (29) were taken as a reference, which included the use of pooled QCs and blanks (for carryover assessment). Additional information regarding this measurement is provided in the supplementary material (Table S4, Text S9).

**Table 1:**
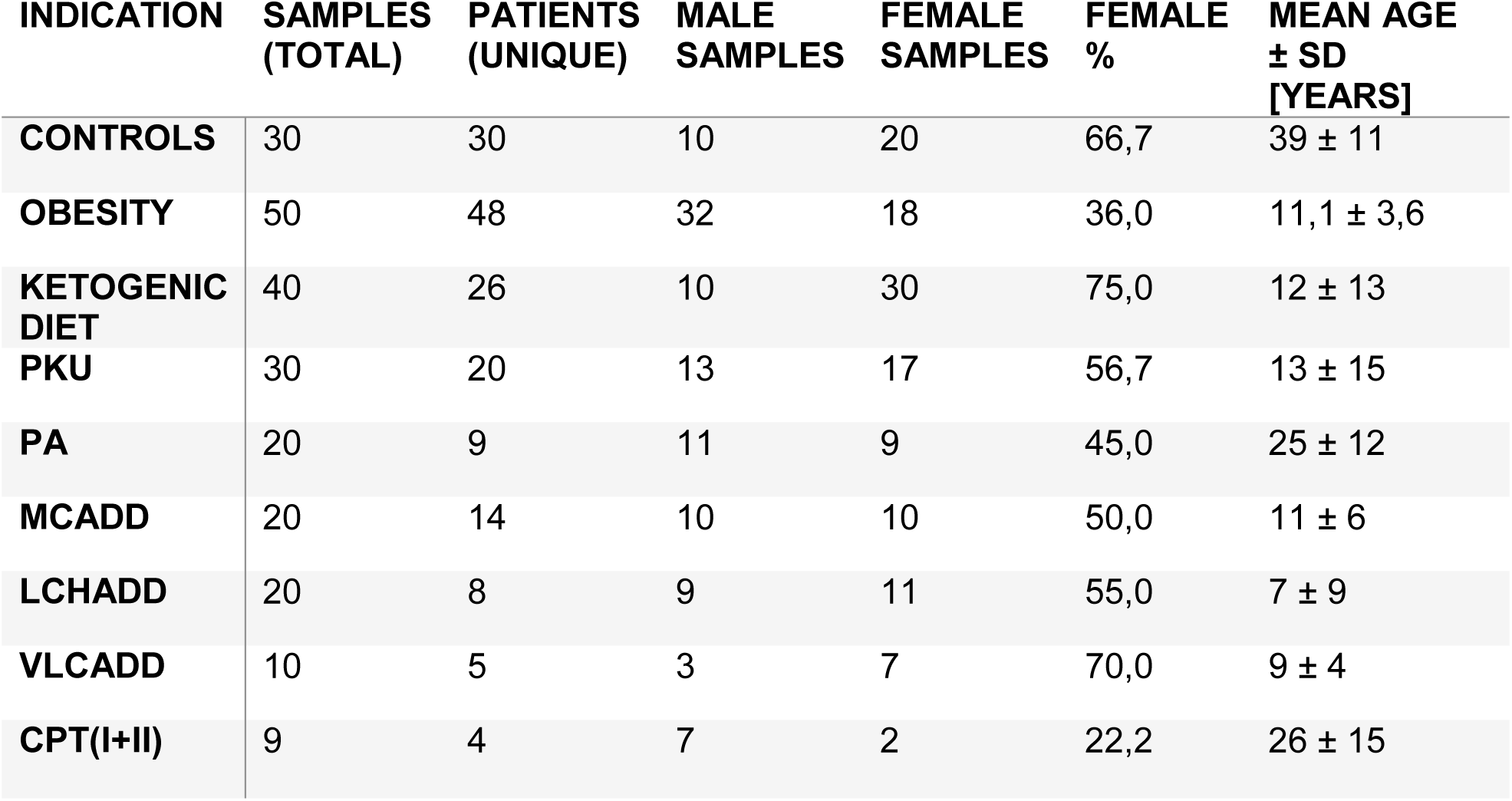
Overview of the residual DBS samples utilized.

### Data analysis

Processing of the raw LC-MS data from the targeted methods was performed with TASQ 2021b (Bruker Daltonics, Bremen, Germany). The recovery of analytes was calculated with the raw peak areas and then by comparing the ISTD-normalized quantified results. Statistical analysis was performed using unpaired t-tests with a correction for multiple comparisons (Holm-Šídák method). P-value < 0.05 were considered significant.

All untargeted metabolomics batches were processed with Metaboscape 2021b (Bruker Daltonics, Bremen, Germany). The T-Rex 3D algorithm for timsTOF in positive ionization mode was applied to automatically extract data and assign the MS2 fragment spectra. Masses were recalibrated with sodium formate clusters that were co-injected with every sample. [M+H]^+^ was set as primary ion. Further details are given in the supplementary material (Text S8, S9).

### Data visualization

Data was visualized with GraphPad Prism (Version 10.2.3), RStudio and Microsoft Excel. MetaboAnalyst 6.0 (30) was used to analyze untargeted metabolomics data. Workflow illustrations were designed with BioRender (Toronto, Canada).

## Results

### Filter paper-induced matrix effects in amino acids and acyl carnitine analyses

To investigate the magnitude of matrix effects introduced by filter paper, we designed an experimental setup (Fig. 1A) in which identical liquid samples are either (i) extracted directly or (ii) applied to filter paper before being subjected to metabolite extraction. Using this approach, acylcarnitines and amino acids were quantified by established LC-MS/MS methods, and the results were compared on the level of raw peak areas as well as ISTD-quantified concentrations. Figure 1 depicts the results for serum as sample matrix, artificially spiked to mimic normal (level 1) and high (level 2) metabolite concentrations.

Most acylcarnitines demonstrated a peak area recovery exceeding 75% (Fig. 1B), including free carnitine (C0) at 92.1%, acetylcarnitine (C2) at 86.6%, butyrylcarnitine (C4) at 87.0% and hexanoylcarnitine (C6) at 92.1%. Recovery rates for quantified concentrations approached close to 100% in most instances. In contrast, dicarboxylic acylcarnitines (DCs), such as C3DC, C4DC and C5DC, exhibited pronounced matrix effects, with peak area recoveries of only 2.9%, 6.7%, and 3.9%, respectively. Here, ISTD-normalization was effective only for C4DC, but with low accuracy. A highly similar pattern was observed in level 2 serum (Fig. 1C), suggesting that the effects are not primarily dependent on the underlying analyte concentration. Statistical comparison between peak area and concentration recoveries further highlighted the substance class specific loss of DCs that potentially arises from an increased interaction with the filter paper (Fig. 1D). Regarding amino acids, the majority of compounds demonstrated satisfactory recoveries, generally close to 100%. Notable exceptions were cystine, cystathionine and asparagine, which showed markedly low peak area recoveries of 14.8%, 7.3%, and 60.9%, respectively, in level 1 serum (Fig. 1E). As a result, the quantification of these metabolites utilizing ISTD-normalization was adversely affected, displaying high variability — an effect that was even more pronounced in level 2 serum (Fig. 1F). Furthermore, aspartate, serine and 2- aminobutyric acid (2-ABA) exhibited a reduced precision compared to other amino acids in both serum levels (Fig. 1E, 1F).

Next, we utilized the same experimental setup to characterize the matrix effects with authentic plasma and whole blood as sample matrices. Lithium heparin whole blood was obtained from a healthy donor and plasma was isolated through centrifugation. In line with previous results obtained from serum, filter paper-induced matrix effects were prominent for DCs in plasma and whole blood (i.e. C3DC, C4DC, C5DC and C6DC; Fig 2, left). For the remaining set of acylcarnitines, the ISTD-normalization was again able to compensate for reduced peak area recoveries. Regarding amino acids, matrix effects affected the peak areas particularly of glutamate (47.3%) and aspartate (48.9%) (Fig. 2, right). Cystine was only detectable in plasma, with a mean peak area recovery of 68% and accompanied by a substantial variability (32%) strongly indicating that it is particularly susceptible to matrix effects. Other amino acids previously observed in serum with reduced recovery and concentrations below the LOD were cystathionine and hydroxyproline. General matrix effects trends, such as the selective loss of specific metabolite classes (e.g. DCs), consistently aligned with previous findings in serum (Fig. 1).

**Figure 2:**
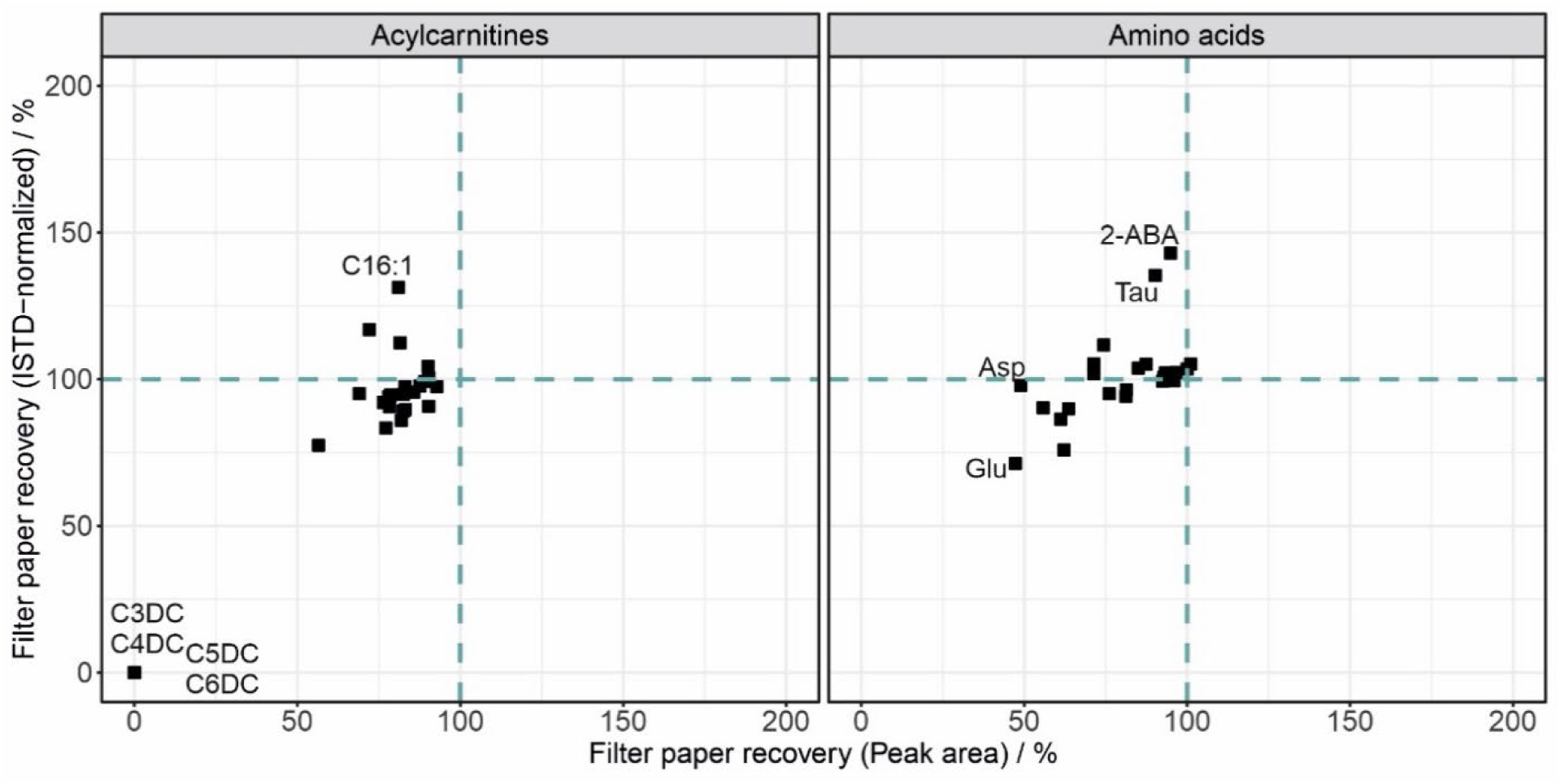
Matrix effects occur for specific compounds in filter paper, as shown for lithium- heparin whole blood. Point plots compare the peak area against the ISTD-normalized recovery. (Left) Acylcarnitines. (Right) Amino acids. Data are presented as mean of five technical replicates (n = 5). Comprehensive results for whole blood and plasma are given in the supplementary material (File 1 and Fig. S3).

Despite the fact that matrix effects were generally compensated using ISTDs, quantitative deviations from filter paper eventually persist for metabolites that lack their own dedicated ISTD. Such discrepancies can even arise when the analyte and ISTD are chemically similar. For example, C16:1 demonstrated an ISTD-normalized recovery of 131% when quantified using the structurally related C16:0-ISTD. Similar overestimations were observed for certain amino acids, including 2-ABA and taurine.

### Filter paper matrix effects in untargeted metabolomics

As anticipated, our targeted mass spectrometry results demonstrated that ISTD correction has the potential to substantially compensate for matrix effects. However, in untargeted metabolomics, individual standards are not available for most metabolites, and even achieving comprehensive coverage at the level of substance classes using commercially available labelled compounds is near impossible. Therefore, we investigated how filter paper-associated matrix effects affect untargeted metabolomic measurements. We conducted an LC-MS/MS analysis with the previously obtained plasma and whole blood extracts. Analytes were separated by Hydrophilic Interaction Liquid Chromatography (HILIC) and detected on a timsTOF Pro mass spectrometer (Bruker Daltonics, Bremen, Germany) with a standardized 3D metabolomics detection method in a scan range from 20-1,300 m/z (see Material and Methods).

Raw data analysis of plasma and whole blood samples was conducted with Metaboscape (Bruker Daltonics, Bremen, Germany) and generated 21,742 and 16,777 features with 202 and 204 annotated compounds, respectively. 27 of these annotations corresponded to ISTDs added to the samples.

PCA analysis of the plasma dataset revealed principle component 1 (PC1) explained 47.4% of the total variance and strongly separated the liquid from the filter paper extraction (Fig. 3A). This indicated that most of the variance in this untargeted metabolomics dataset could be explained by matrix effects introduced by the filter paper. In total 37 features were overrepresented in liquid extraction conditions by a minimum 1.5-fold change (FC) (Fig. 3B). The whole blood metabolomics dataset also showed highly similar results, with PC1 explaining 62.2% of the variance in an extraction-condition dependent manner, with 45 upregulated and five downregulated compounds (Fig. 3D).

**Figure 3:**
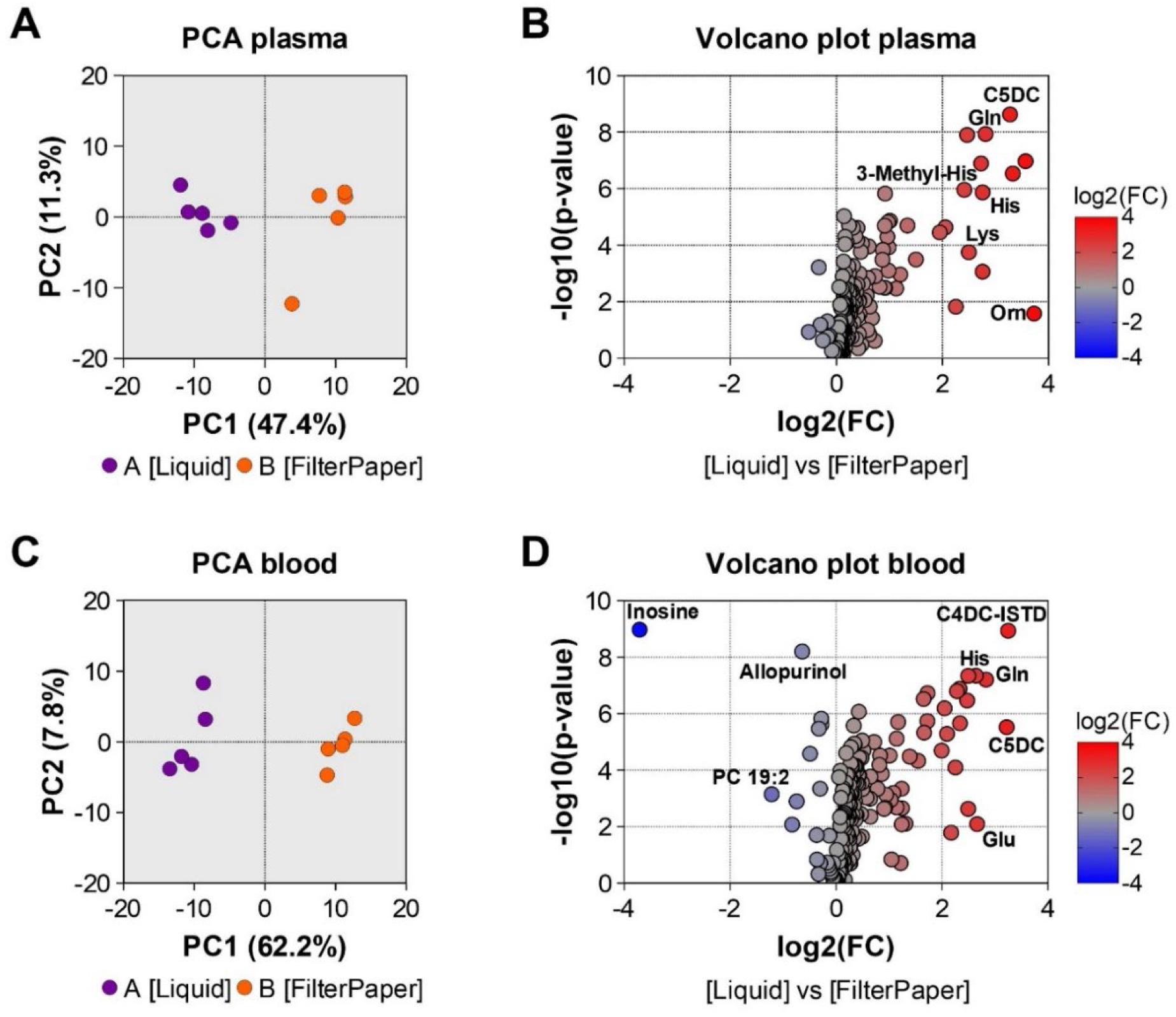
Untargeted metabolic profiles differ between liquid plasma / blood and their respective filter paper analogue. (A) A PCA from plasma separates both extractions. (B) A volcano plot reveals several upregulated compounds in the liquid extraction, including amino acids like histidine and ornithine as well as the previously identified C5DC acylcarnitine compound. (C) A PCA from blood separates both extractions as well. (D) A volcano plot from blood reveals several up- and downregulated features. A full list of identified compounds is given in supplementary material (File 2).

A general observation was that most of the significantly altered metabolites exhibited losses upon application to filter paper, with almost none showing enrichment. One of the few exceptions was inosine in the blood sample matrix. Consistent with findings from the targeted approach, dicarboxylic acylcarnitine species (C5DC) were found to be highly affected by matrix effects, along with several charged and polar amino acids (e.g. Gln, His, Lys). Importantly, a large fraction of metabolites (82% in plasma and 76% in whole blood) retained stable mass spectrometric signals following filter paper application, indicating minimal impact of this step for the respective subset of compounds.

### Filter paper as sample matrix for untargeted analysis of inborn errors in metabolism

In light of the observed matrix effects, we next evaluated the suitability of untargeted metabolomics for distinguishing biomarker profiles within a patient cohort exhibiting diverse metabolic perturbations. A total of 229 unique dried blood spot samples were obtained from 30 healthy controls, as well as from diagnostic residual samples of individuals with clinical indications such as obesity, ketogenic diet, PKU, PA, MCADD, LCHADD, VLCADD and the CPT1/CPT2 deficiencies (details given in Materials and Methods). Samples were extracted and subjected to untargeted metabolomics analysis, which led to 195 identified compounds spanning 7 superclasses and 23 classes (Fig. 4).

**Figure 4:**
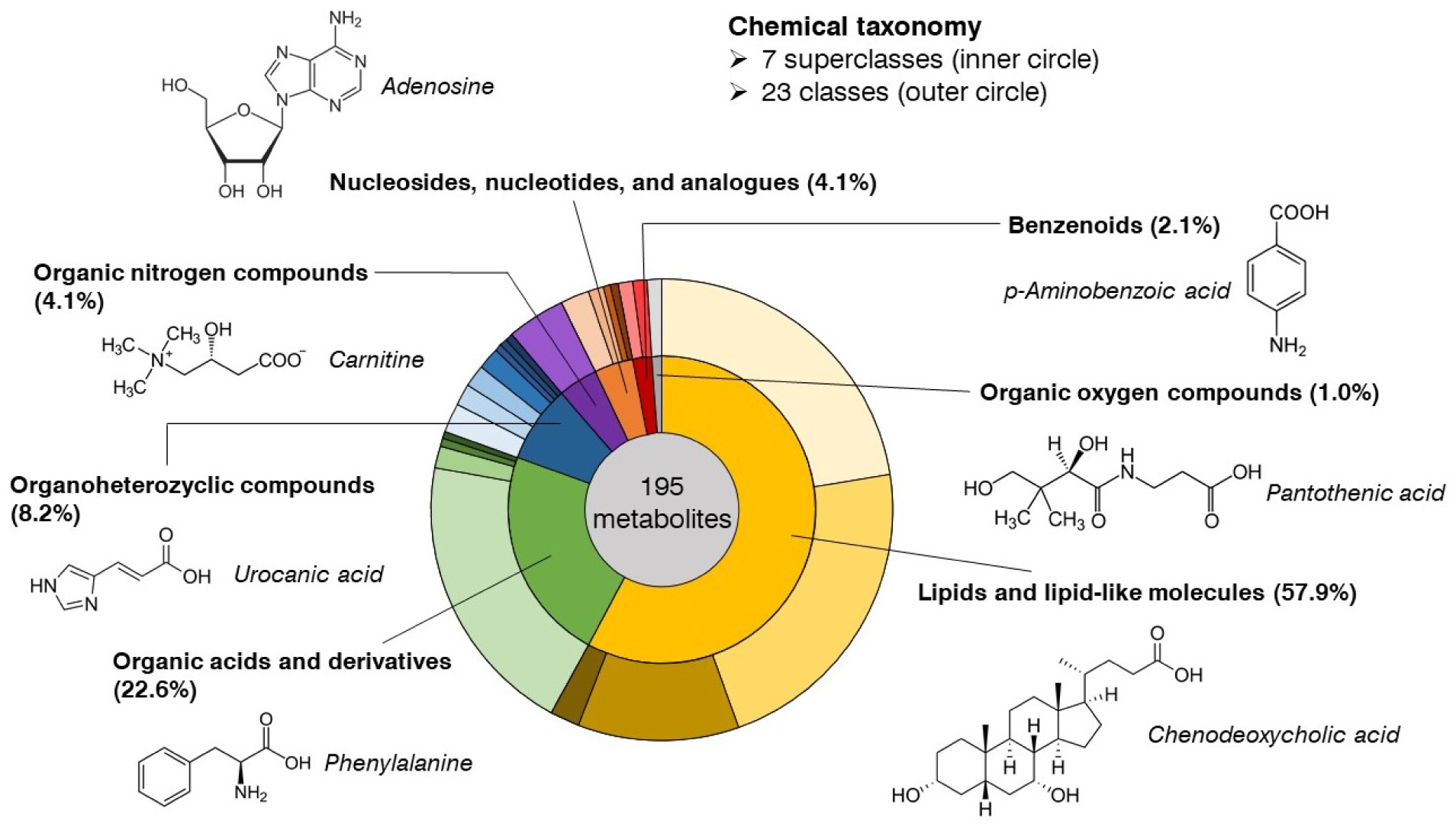
Qualitative analysis of identified metabolites with untargeted metabolomics from DBS extracts in HILIC mode. The compounds belong to seven different superclasses, with a dominant fraction of lipids and lipid-like molecules followed by organic acids and derivatives. These superclasses then converge into 23 different classes. A detailed list of identified metabolites, superclasses and classes is given in the supplementary material (File 3).

The metabolic diversity of the cohort with respect to the underlying disturbances is reflected in a PCA where only a relatively small portion of the variance (8.13%) is captured by principal component 1. Nonetheless, influences from specific conditions such as LCHADD and ketogenic diet are still discernible at this level (Fig. 5A). Next, we used this untargeted metabolomics dataset to evaluate whether the different medical conditions could be classified based on known biomarkers, despite the previously described matrix effects and in the absence of internal standard correction. Upon using phenylalanine (Fig. 5B), C3 (Fig. 5C), C4OH (Fig. 5D), C8 (Fig. 5E), C14:1 (Fig. 5F) and C16OH (Fig. 5H), the respective indications of PKU, PA, ketogenic diet, MCADD, LCHADD and VLCADD are already reliably separated from other DBS samples. However, C16 (Fig. 5G) levels from LCHADD patients are within the normal range of the cohort, indicating a rapid metabolic conversion of C16 into C16OH. Notably, a subset of DBS samples (∼ 2 for PKU, ∼ 5 for MCADD, ∼ 8 for ketogenic diet) remains indistinguishable from the rest, which is less surprising considering that the metabolic state of some (younger) patients is medically treated or at least permanently controlled, in order to minimize any severe IMD-related disease symptoms.

**Figure 5:**
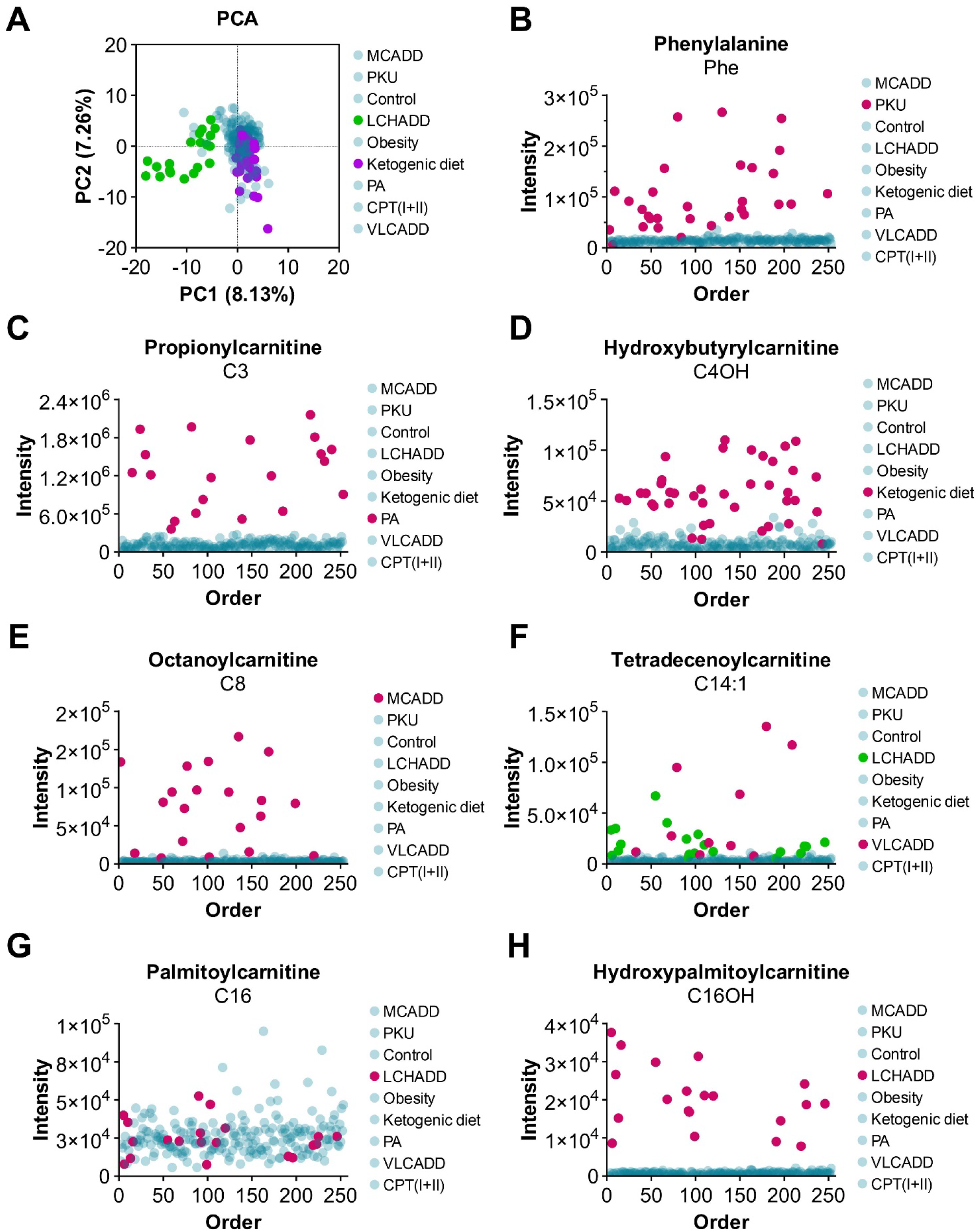
HILIC-MS untargeted metabolomics can serve as a tool to differentiate multiple indications from a DBS cohort. (A) A PCA analysis reveals that LCHADD and ketogenic diet are separated to a certain degree from the rest of the cohort. (B) – (H) Respective intensity responses from amino acid and acylcarnitine analytes were plotted against the injection order of the DBS samples to highlight separations from different clinical indications.

In clinical practice, the variability of biomarker concentrations is often compensated by calculating ratios that serve as surrogate diagnostic markers for IMDs. Following this approach and using the untargeted metabolomics intensities as underlying data source successfully distinguishes patients diagnosed with PKU (Fig. 6A), PA (Fig. 6B), MCADD (Fig. 6C), LCHADD (Fig. 6D) and to a certain degree VLCADD (Fig. 6E) from the cohort. The (C16+C18:1)/C2 ratio, which can be indicative of CPTII deficiency, showed the lowest discriminative power (Fig. 6F), which is likely due to the influence of the dietary treatment – e.g. limitation of long-chain fatty acids in LCHADD and VLCADD patients – associated with several of the included conditions.

**Figure 6:**
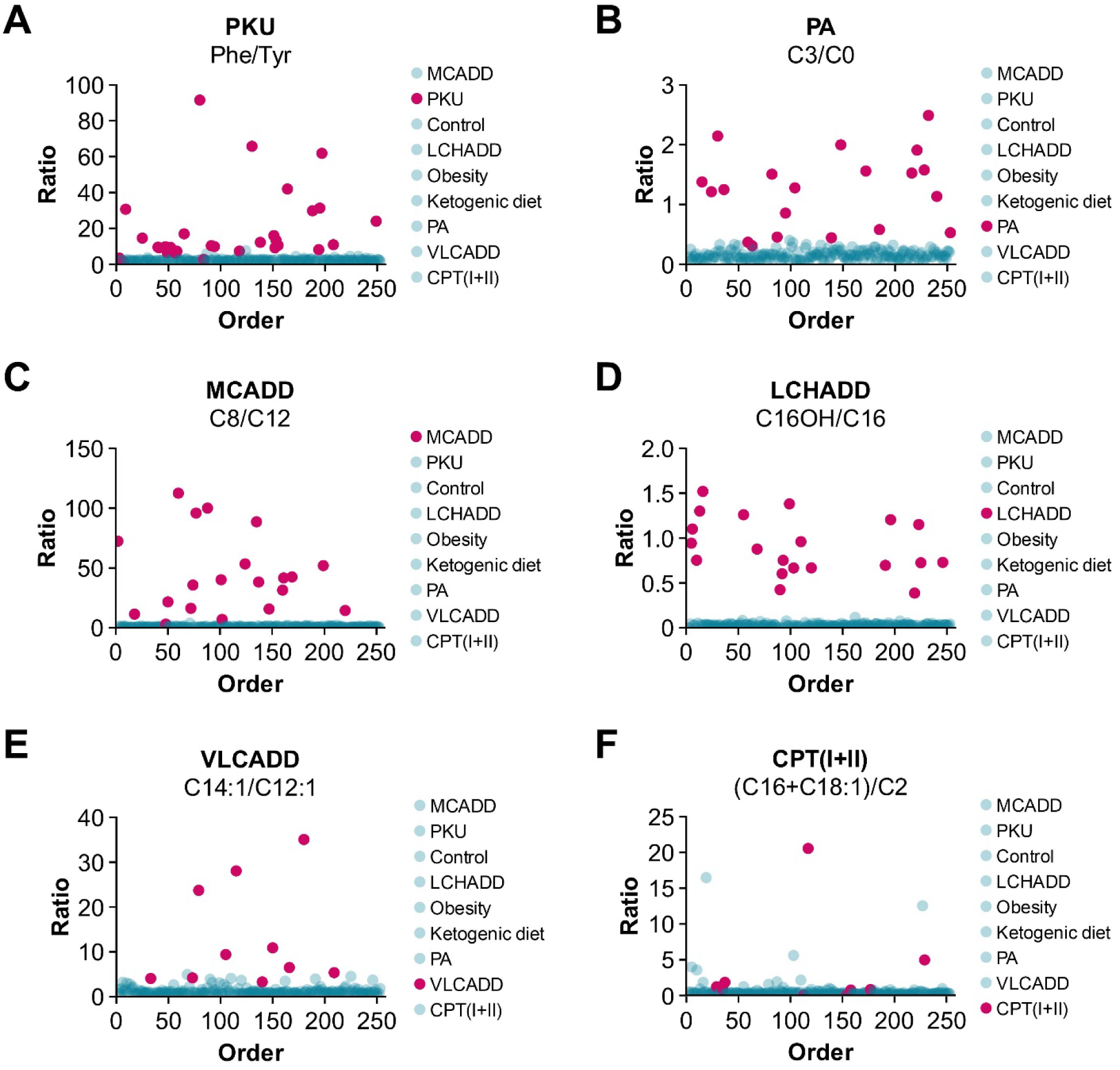
HILIC-MS derived intensity responses from metabolites were used to generate IMD-related ratios. (A) Phe/Tyr ratio for PKU. (B) C3/C0 ratio for PA. (C) C8/C12 ratio for MCADD. (D) C16OH/C16 ratio for LCHADD. (E) C14:1/C12:1 ratio for VLCADD. (F) (C16+C18:1)/C2 ratio for the grouped CPT(I+II) disorders.

As the diversity of the untargeted metabolic profiling dataset was too complex for generating a discriminating PCA we conducted a random-forest (RF) machine learning analysis (Fig. 7A). In a first step we optimized RF parameters such as the number of trees (which yielded stable results at >2000) and the number of predictors (supplementary Fig. S4). The best model had an overall out-of-bag (OOB) error of 7.0%. We found only small class errors for most groups, i.e., 0% for obesity, controls and LCHADD, 2.5% for ketogenic diet, 3.3% for PKU, 5% for PA, and 10% for MCADD (compare Fig. 7B). In contrast, an accurate classification of VLCADD and CPT(I+II) deficiencies based on untargeted metabolomics data was challenging for the model and was associated with an OOB error of 50% and 66%, respectively (Fig. 7C). Two factors that may explain this observation: i) VLCADD and CPT(I+II) have the smallest sample sizes (Table 1), and ii) these conditions are potentially associated with only a limited set of highly informative metabolite features. To assess whether the inclusion of additional information could enhance the selectivity of the model, we enriched the dataset with 15 metabolite ratios commonly used in the diagnosis and monitoring of these conditions, and repeated the analysis. The ratios were calculated directly from the intensities in the untargeted metabolomics dataset. The extended model showed improved results specifically for the VLCADD group but had little influence on the other groups and provided no additional benefit for CPT(I+II) deficiencies (Fig. 7D). Overall, the OOB error decreased slightly from 7.0% to 6.1%, indicating that artificially expanding untargeted metabolomics dataset by constructing ratios can have the potential to improve the performance of RF. In this context, an analysis of the top 10 most important features identified by both RF models provides meaningful insights into the variables driving classification (Fig. 7E-F). The RF model based on the untargeted dataset is strongly driven by a set of clinically well-known biomarkers including several acylcarnitines such as hydroxybutyrylcarnitine (C4OH) and octanoylcarnitine (C8) as well as the amino acids phenylalanine and tyrosine. Interestingly, methylmalonylcarnitine (C4DC), a metabolite that in our previous analysis showed to be strongly affected by filter paper-caused matrix effects (compare Fig. 1B, 1C), also was highly important for the RF model. In the ratio- enriched dataset the importance of many of these metabolites is outperformed by the ratios that derive from them (Fig. 7F), demonstrating that these ratios offer added value beyond the directly measured biomarkers themselves.

**Figure 7:**
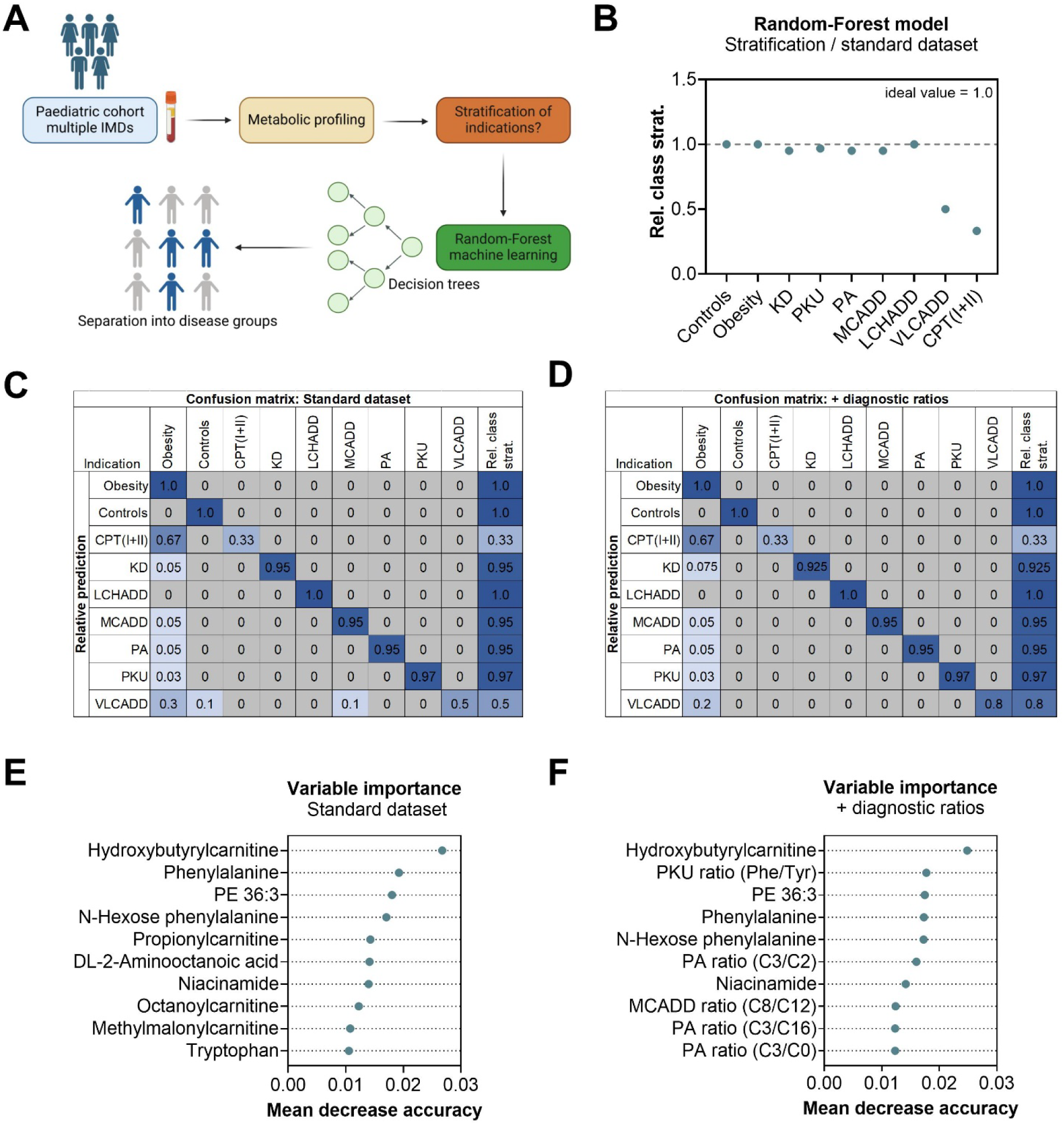
Application of a Random-Forest (RF) machine learning analysis onto a clinical DBS cohort. (A) The schematic workflow depicts the flow starting from a clinical paediatric cohort over the metabolic profiling of DBS samples to the application of the RF analysis. (B) The RF model shows excellent stratification results for seven indications. (C) The complete confusion matrix performed with the standard dataset. (D) Same as C, but with additional diagnostic ratios. VLCADD shows strongly improved results. (E) The ten most important features for the RF model in C are ranked by their mean decrease accuracy. (F) Same as E, but related to the dataset with diagnostic ratios.

N-Hexose phenylalanine strongly contributed to the stratification of PKU samples in our study and was ranked at a high 4^th^ place within the variable importance (Fig. 7E and supplementary Fig. S5). This compound has already been reported earlier as an potential secondary PKU biomarker in plasma (31) and was found by us in an unsupervised manner also when using dried blood spots as sample matrix.

## Discussion

In this study, we investigate matrix effects induced by filter paper that is used for DBS sampling. This sample material represents a well-established method for the collection, transport and storage of specimens in clinical practice and is integrated in numerous workflows (32, 33), including newborn screening programs, and has remained largely unchanged over the decades (11). The evolution of analytical technologies employed, in particular mass spectrometric biomarker analysis, allows concomitant analysis of numerous metabolites (34) and may reveal important limitations of filter paper as sampling matrix for specific metabolites. Here, we assessed the performance and potential limitations of filter paper in an untargeted metabolomics workflow (Fig. 3).

The rationale of focusing on DBS is based on practical realities. Emerging technologies aim to improve the handling of clinical samples (35–37), and robotic systems reduced the analytical variation when compared to manual sample preparation (38). However, the implementation of such novel sampling systems is often hindered by challenging factors such as costs, limited availability of trained personnel, or special logistics. As a result, the real-world decision is often not between DBS and a better, more advanced alternative, but rather between DBS and no sample at all. Thereby, it is of critical importance to maximize the scientific and clinical value derived from this simple, cost-effective, and widely accessible sample type.

In targeted methods, matrix effects caused by filter paper can largely be compensated by utilizing matched internal standards (Fig. 1B-C, 1E-F). As shown by us (Fig. 3) and others (29, 39), for untargeted metabolomics this is much more difficult because of not only the limited availability of stable isotope-labelled standards but also the extensive chemical diversity of the compounds of interest (Fig. 4). Even within a single class – such as amino acids – matrix effects (Wang et al. in 2013 (40)) as well as ionization properties can vary considerably (Fig. 5C-H) (41). This renders standardization across a broader range of metabolites in untargeted mass spectrometric analyses challenging.

A considerable portion of the matrix effects reported in this study likely arise from the high surface area of filter paper and the polar interactions with its cellulose structure (42). The drying process in DBS further concentrates metabolites at this surface, enhancing matrix interactions. It is therefore not surprising that we observed pronounced matrix effects particularly for dicarboxylic acylcarnitines, with peak area recovery losses of more than 90% for C3DC, C4DC, C5DC, and C6DC (Fig. 1B - D, Fig. 2 left). These effects were independent of both, concentration and biological sample type, suggesting a chemically driven mechanism. Dicarboxylic acylcarnitines are zwitterionic molecules with two carboxyl groups and a relatively low molecular weight, which makes them especially prone to strong interactions with the filter paper. One potential strategy to mitigate such effects is to optimize extraction protocols tailored to the most affected compound classes. Indeed, various studies have used a diverse spectrum of extraction solvents ranging from pure methanol or acetonitrile or their mixture to more complex mixtures (43–45). For untargeted analytical strategies, a compromise has to be found that accommodates a broad range of biomarkers, with the effect that matrix effects and reduced extraction efficiencies for specific compound classes represent unavoidable trade-offs.

More than 1500 IMDs have been identified so far (18), and there is also a wide range of polygenic or multifactorial metabolic disorders. Targeted mass spectrometric methods are currently only able to provide meaningful diagnostic information for a very limited subset of these conditions (46). This underscores the significant potential of broader analytical strategies capable of reliably capturing a spectrum of metabolite classes, with untargeted metabolomics representing one such promising approach. In order to maximise the number of reliable features, a careful balance between achieving accurate compound identification and maintaining the exploratory nature of the approach must be found. The untargeted approach utilized in this study identified 195 high confidence compounds in positive ionisation mode, aligning well with similar, previously reported studies (21, 47). However, without the possibility for reliable absolute quantification it is difficult to perform quantitative comparisons with previously established reference values. Untargeted datasets not only have limited capacity to correct for matrix effects, they often also require statistical batch correction to achieve longitudinal comparability (48, 49). Consequently, the clinical interpretation of individual diagnostic samples becomes challenging when they cannot be analyzed in the context of a larger cohort. This is particularly relevant in clinical practice as diagnostic decisions often need to be made on the basis of single patient samples.

We thus investigate the applicability of an untargeted metabolomics workflow in a well defined clinical DBS cohort of metabolic conditions, including several IMDs (i.e. PKU, PA, MCADD, LCHADD and VLCADD). Notably, the raw intensity values of known biomarkers proved to be highly indicative of the underlying disease even in the absence of absolute quantification and the potential presence of matrix effects (Fig. 5). The diagnostic stability of the signals can be further enhanced by using raw intensity values to reconstruct established clinical ratios (Fig. 6). The effectiveness of this approach depends on the quality of the ratios and their susceptibility to external influences. In our study, this could be observed for the (C16+C18:1)/C2 ratio, which is primarily indicative of CPT II deficiency (Fig. 6F), but is also readily modulated by nutritional intervention therapies for other IMDs. In general, our data indicate that achieving a clear differentiation of CPTI/II deficiencies based on untargeted metabolomics data is more difficult compared to the other selected IMDs (compare Fig. 5 and 6F). This aligns with previous observations that CPT II deficiency may present with normal acylcarnitine profiles in DBS, and with existing recommendations to perform analysis in plasma samples instead (41).

Despite the above described inherent challenges of DBS-based untargeted metabolomics data sets, we were able to achieve a clear differentiation between distinct metabolic disease by means of a random forest model (Fig. 7), an approach that also performed well in a range of other metabolite datasets (50–52). The integration of computational methods to improve the manual diagnosis of IMDs has recently demonstrated accurate and reproducible identification of 16 IMDs (53). In our study, an analysis of the most important features driving successful classification revealed that many of these biomarkers overlap with metabolites already established in clinical practice through targeted assays (54–56). At the same time, the untargeted approach also offers the opportunity to identify novel or previously underexplored biomarkers, potentially expanding the current diagnostic repertoire. In this line, we observed the accumulation of N-Hexose Phe conjugates in DBS of PKU patients (Fig. 7 E-F). Further literature research revealed that this biomarker had already previously been described in patient plasma (31) and was subsequently subjected to more detailed structural analysis (57).

In summary, filter paper used DBS sampling increased the risk of matrix effects, which are often specific to certain compound classes. These effects can typically be corrected by choosing suitable internal standards unless the signal suppression lowers analyte intensities into a suboptimal signal-to-noise range. Nonetheless, even without correction, untargeted metabolomics datasets contain valuable disease-relevant information that can be extracted using appropriate analytical strategies. Combining untargeted approaches with established clinical ratios may offer a powerful strategy to balance broad metabolic coverage with diagnostic relevance and at the same time maintaining the potential for discovery.

## Author contributions

J.C. and M.A.K. designed the research; J.Z. and D.K. contributed with medicinal input; J.C. performed research; J.C. and J.K. contributed new reagents/analytic tools; J.C. and V.J. analyzed data; J.C. and M.A.K. wrote the paper with contributions from all authors.

## Acknowledgements

The authors thank all participants of the metabolomics study.

## Ethics declaration

The data used in the study were generated in the course of standard diagnostic processes with full informed consent from the investigated individuals. For study purposes, all data were pseudonymized by removing individual personal, clinical or other information, and fully anonymized after completion of the comparative analyses. The analyses were approved by the Clinical Research Ethics Board of the Medical University Innsbruck (votum 1320/2020 and 1163/2024).

## Data availability

The untargeted metabolomics data (from the DBS study cohort) with identified compounds and intensities was made publicly available under a CC-BY-NC-4.0 license on Mendeley Data (DOI: 10.17632/vc8wnf62jg.1).

