## Supplemental Material for "Matrix effects influence biochemical signatures and metabolite quantification in dried blood spots"

**Table S1: Chromatographic gradient profile for acylcarnitine method.**

| **Time** | **Flow rate / ml*min$^{-1}$** | **Mobile Phase A / %** | **Mobile Phase B / %** | **Gradient shape** |
|---|---|---|---|---|
| 0.00 | 0.400 | 2.0 | 98.0 | Linear |

| | | | | |
|---|---|---|---|---|
| 0.50 | 0.400 | 2.0 | 98.0 | Linear |
| 7.00 | 0.400 | 7.0 | 93.0 | Linear |
| 9.00 | 0.400 | 40.0 | 60.0 | Linear |
| 9.50 | 0.500 | 60.0 | 40.0 | Linear |
| 10.50 | 0.500 | 60.0 | 40.0 | Linear |
| 11.50 | 0.400 | 2.0 | 98.0 | Linear |
| 12.00 | 0.400 | 2.0 | 98.0 | Linear |
| Equilibration time: 0.00 min | | | | |

**Table S2: Chromatographic gradient profile for amino acid method.**

| Time | Flow rate / $ml*min^{-1}$ | Mobile Phase A / % | Mobile Phase B / % | Gradient shape |
|---|---|---|---|---|
| 0.00 | 0.400 | 2.0 | 98.0 | Linear |
| 3.00 | 0.400 | 2.0 | 98.0 | Linear |
| 6.00 | 0.400 | 7.0 | 93.0 | Linear |
| 8.00 | 0.500 | 55.0 | 45.0 | Linear |
| 10.00 | 0.500 | 55.0 | 45.0 | Linear |
| 10.30 | 0.400 | 2.0 | 98.0 | Linear |
| 11.00 | 0.400 | 2.0 | 98.0 | Linear |
| Equilibration time: 1.00 min | | | | |

**Table S3: timsTOF-MS parameters for ESI-source methods.**

| Method | Acylcarnitine method | Amino acid method | Standard metabolomics method |
|---|---|---|---|
| Instrument | timsTOF Pro (Bruker Daltonics, Bremen, DE) | | |
| Source | ESI | | |
| Ionization mode | positive | | |
| Tims | disabled | | |
| Calibration | 5 mM Sodium formiate solution in 90% IPA | | |
| Calibration mode | Enhanced quadratic | quadratic | Enhanced quadratic |
| Scan range | 50 – 500 m/z | 50 – 350 m/z | 20 – 1300 m/z |
| Spectra Rate | 5.00 Hz | 4.00 Hz | 12.00 Hz |

| | | | |
|---|---|---|---|
| End plate offset | 500 V | 500 V | 500 V |
| Capillary voltage | 4500 V | 4500 V | 4500 V |
| Nebulizer | 2.5 bar | 2.5 bar | 2.2 bar |
| Dry gas | 10.0 l/min | 10.0 l/min | 10.0 l/min |
| Dry temp. | 250°C | 250°C | 220°C |
| MS Peak Detection | 100 cts. (max. intensity) | 100 cts. (max. intensity) | 25 cts. (max. intensity) |
| Tuning parameters | | | |
| Deflection Delta 1 | 60.0 V | 60.0 V | 60.0 V |
| Funnel 1 RF | 150.0 Vpp | 90.0 Vpp | 200.0 Vpp |
| Funnel 2 RF | 200.0 Vpp | 250.0 Vpp | 200.0 Vpp |
| isCID Energy | 0.0 eV | 0.0 eV | 0.0 eV |
| Multipole RF | 150.0 Vpp | 150.0 Vpp | 60.0 Vpp |
| Ion Energy | 3.0 eV | 3.0 eV | 5.0 eV |
| Low Mass | 50.00 m/z | 50.00 m/z | 60.00 m/z |
| Collision Energy | 5.0 eV | 5.0 eV | 7.0 eV |
| Pre Pulse Storage | 5.0 µs | 5.0 µs | 5.0 µs |
| Stepping Mode | Advanced | Basic | Basic |
| Stepping Collision RF | 150.0; 300.0; 450.0; 600.0 (equivalent time) | 300.0 Vpp to 600.0 Vpp (Timing: 60/40%) | 200.0 Vpp to 700.0 Vpp (Timing: 50/50%) |
| Stepping Transfer Time | 20.0; 35.0; 50.0; 65.0 | 20.0 to 35.0 µs | 20.0 to 80.0 µs |
| Collision Energy | 100; 100; 200; 200% | 100 to 100% | 100 to 250% |
| MS/MS mode | bbCID; 50.0 eV | bbCID; 20.0 eV | Auto MS/MS |

**Table S4: timsTOF-MS parameters for VIP-HESI source method.**

| **Method** | **Optimized untargeted metabolomics method (used for 229 DBS cohort)** |
|---|---|
| Instrument | timsTOF Pro (Bruker Daltonics, Bremen, DE) |
| Source | VIP-HESI |
| Ionization mode | positive |
| Tims | disabled |
| Calibration | 5 mM Sodium formiate solution in 90% IPA |
| Calibration mode | Enhanced quadratic |
| Scan range | 20 – 1300 m/z |
| Spectra Rate | 4.00 Hz |
| End plate offset | 500 V |

| | |
|---|---|
| Capillary voltage | 4500 V |
| Nebulizer | 3.0 bar |
| Dry gas | 10.0 l/min |
| Dry temp. | 260°C |
| Probe Gas Temp | 400 °C |
| Probe Gas | 5.0 l/min |
| Exhaust | activated |
| MS Peak Detection | 100 cts. (max. intensity) |
| Tuning parameters | |
| Deflection Delta 1 | 60.0 V |
| Funnel 1 RF | 150.0 Vpp |
| Funnel 2 RF | 225.0 Vpp |
| isCID Energy | 0.0 eV |
| Multipole RF | 100.0 Vpp |
| Ion Energy | 5.0 eV |
| Low Mass | 50.00 m/z |
| Collision Energy | 5.0 eV |
| Pre Pulse Storage | 5.0 µs |
| Stepping Mode | Basic |
| Stepping Collision RF | 300.0 to 700.0 Vpp (Timing: 50/50%) |
| Stepping Transfer Time | 20.0 to 80.0 µs |
| Collision Energy | 100 to 100% |
| MS/MS mode | Auto MS/MS (Cycle Time: 0.5 s), active exclusion after 3 spectra, threshhold 350 cts, smart exclusion 5x |

### Text S5: Acylcarnitine method details.

Preparation of internal standards:

The internal standards (NSK-B-1 = AC-ISTD-1 and NSK-B-G1 = AC-ISTD-2) are prepared according to the vendors instructions. After 1.00 mL of $H_2O$ (LC-MS grade) are added, closed and vortexed, 10 µL from this solution are aliquoted to fresh ND-10 vials. The vials are evaporated under N2-flow, closed and stored at -20°C until day of preparation.

Preparation of samples:

1. To a vial of AC-ISTD-1, 1500 µL of ACN:MeOH (3:1 v/v) are added.
2. To a vial of AC-ISTD-2, 1600 µL of ACN:MeOH (3:1 v/v) are added.
3. Both ISTDs are vortexed.
4. 20 µL of plasma or 1 DBS spot (6.00 mm) are placed in a 1.5 mL screw-cap vial.
5. 100 µL are added from AC-ISTD-1, then 100 µL from AC-ISTD-2.
6. To this solution, 300 µL of extraction solvent (ACN:MeOH (3:1 v/v)) are added.
7. A small spatula tip (~ 100 mg) of glass beads are added.
8. Vials are then shaken for 2.5 min at 20 Hz using a MM400 mixer mill.
9. Vials are then placed in an ultrasonic bath for 5 minutes.
10. Vials are vortexed for 10 seconds.
11. Then, the vials are centrifuged at 31150 x g (4°C) for 10 minutes.
12. The supernatant is transferred to an autosampler vial and evaporated under N2-flow.
13. 100 µL of extraction solvent (ACN:MeOH (3:1 v/v)) are added again. From this solution, 10 µL are injected into the column with the described LC-MS method for acylcarnitines.

This sample preparation was used for the method validation and is used for real diagnostic samples in our lab. For the samples in the paper, an adjustment had to be made, in order to not overload the pre-placed filter paper spot with liquid sample. In the paper, 11 µL of liquid sample was used and 11 µL of liquid sample were placed directly onto a empty, pre-placed filter paper spot. The rest of the above shown preparation protocol remained the same.

Accuracy: The concentration accuracy was tested out by measuring two lyophilized human serum samples from Erndim were reference values were given by a peer laboratory.

Precision: To account for realistic batch conditions, we chose to determine the precision by using a QC sample (pool of 82 plasma) that was injected 11 times, while measuring 8 samples in between covering a total of 24 hours. For determining the Coefficient of Variation (CV), we selected a 99.9% cutoff of total acylcarnitine content resulting in n = 21 and calculated a median CV of 4.77 %.

Linear Dynamic Range: The linear range was first determined by a dilution series of twelve acylcarnitines. All standards, except Glutarylcarnitine, display a similar behavior with a linear response between approximately 0.005 pmol and 5-10 pmol, thus approximately 3 magnitudes of power, before reaching a signal plateau. This includes Hydroxyacylcarnitines as well as stable isotope labeled N-methyl-$D_3$ acylcarnitines. The only carboxy-acylcarnitine Glutarylcarnitine (C5DC), however, displays a limited linear response between 0.1 pmol and 5 pmol. For free carnitine and ten diagnostically relevant acylcarnitines, we determined the linear dynamic range by using a linear regression with a coefficiant of determination ($R^2$) above 0.98. For values within this range, a linear relationship between the Internals Standards and the acylcarnitines can be assumed.

Limit of Detection and Limit of Quantification: A general (mean) LOD and LOQ were determined by diluting 12 acylcarnitine standards until the signal reached the steady-state noise level. The LOD was then calculated by the mean plus three times the standard deviation of the noise, while the LOQ was calculated by the mean plus five times the standard deviation of the noise. Excluding these data points, a quadratic regression representing the acylcarnitine response was created [$y = -0.1224x^2 + 0.7258x + 6.2719$] and used to calculate corresponding peak area ~ molar values. The LOD and LOQ values were 7.25 fmol and 9.6 pmol, respectively.

Robustness: Two commonly used and established extraction procedures for acylcarnitines in plasma were tested and evaluated, one being 100 % methanol, the other 3:1 acetonitrile:methanol. In both protocols, the same total number of acylcarnitines were detected and the extraction had no effect on the elution profile. However, all acylcarnitines extracted with 3:1 acetonitile:methanol had a higher peak area than the other method. Also the addition of amino acid internal standards did not have any effect on intensity values.

Recovery: A pool of plasma was spiked with two concentrations (Level 1 and Level 2) of a defined amount of Acylcarnitines (AC spike mix) either before extraction or added after extraction. All conditions were handled and measured in five independent replicates. ACs were then quantified using the internal standards. The pure plasma and the AC mix were also quantified. Resulting quantified matrix effects can be found in the main manuscript.

### Text S6: Amino acid method details.

Preparation of internal standards: The internal standard (NSK-CAA-1 = AA-ISTD-1) is prepared according to the vendors instructions. After 1.00 mL of $H_2O$ (LC-MS grade) are added, closed and vortexed, 10 µL from this solution are aliquoted to fresh ND-10 vials. The vials are evaporated under N2-flow, closed and stored at -20°C until day of preparation.

Preparation of samples:

1. To one vial of AA-ISTD-1, 1000 µL of MeOH (HPLC-grade) is added.
2. The ISTD is vortexed.
3. 20 µL of plasma or 1 DBS spot (6.00 mm) are placed in a 1.5 mL screw-cap vial.
4. 100 µL are added from AA-ISTD-1.
5. Extraction solvent MeOH is added, the volume depends on the sample type. For plasma, 280 µL are added, while for a DBS sample, 300 µL are added. The total volume is 400 µL in both cases.
6. A small spatula tip (~ 100 mg) of glass beads are added to plasma samples.
7. Vials are then shaken for 2.5 min at 20 Hz using a MM400 mixer mill.
8. Vials are then placed in an ultrasonic bath for 5 minutes.
9. Then, the vials are centrifuged at 31150 x g (4°C) for 3 minutes.
10. The supernatant is transferred to an autosampler vial.
11. From this solution, 10 µL are injected into the column with the described LC-MS method for amino acids.

For the matrix effect recovery samples in the paper, an adjustment was made, in order to not overload the pre-placed filter paper spot with liquid sample: 11 µL of liquid sample was used and placed directly onto a empty, pre-placed filter paper spot. To account for the smaller sample volume, the total extraction solvent volume was adjusted to 200 µL in both preparations, and 5 µL were injected into the column.

<u>Accuracy</u>: The accuracy was again testet out by measuring lyophilized human serum samples from Erndim. The 1-point internal calibration was used for quantification.

<u>Intra- and Interday precision:</u>The intra- and interday precision was directly calculated from the accuracy measurements, as the control samples were injected 5 times per day and the different control samples were prepared on five different days with CV values for most analytes being in the range of 5-15%.

<u>Linearity:</u> A prepared mix of 26 amino acids was diluted in a dilution series from 1500 – 0 µmol/L (Steps: 1500, 1200, 900, 600, 300, 100, 0). 20 µL of the solutions were prepared like a sample and the linearity was calculated by plotting the ratio of analyte/internal standard peak areas against the concentration of the analyte. The $R^2$ was excellent (>0.99) for most amino

acids over the observed range, with exceptions for cystine and analytes without their own respective internal standard.

LOD and LOQ: The limit of detection (LOD) and limit of quantification (LOQ) were calculated by using the signal-to-noise ratio from the linearity measurements, as this value is given in the data processing software TASQ. The LOD and LOQ were defined as:

$$LOD = 3 \times S/N$$

$$LOQ = 10 \times S/N$$

The LOD ranged from 6,6 – 0,01 µmol/L and the LOQ from 22,1 – 0,04 µmol/L. T

Robustness: The robustness was tested out by changing the extraction solvent. The test included a comparison between MeOH, MeOH:H2O (80:20 v/v), ACN:H2O (80:20 v/v), and ACN:MeOH (3:1 v/v). No significant changes in the accuracy were found for the first three solvents, but the ACN:MeOH mixture was found to get less accurate results. The robustness showed that ACN:MeOH (3:1 v/v) might not be usable for amino acid extraction, but small water contents in the extraction solvent do not influence the results.

Recovery from spiking: A plasma sample was spiked with an amino acid standard mix in two levels (Level 1 and Level 2). 3 Replicates were measured with the amino acid method and the concentrations were determined by the 1-point-internal calibration with the internal standards. The recovery was calculated by the ratio of the measured value to the theoretical value from the spike. Excellent results were obtained, with slight exceptions for analytes that did not have their own respective internal standard (2-ABA, Citrulline, Cystathionine, Ornithine, Taurine).

**Table S7: Abbreviations used for compounds in graphs.**

| Name | Abbreviation | Name | Abbreviation |
|---|---|---|---|
| Carnitine | C0 | 2-aminobutyric acid | 2-ABA |
| Acetylcarnitine | C2 | Alanine | Ala |
| Propionylcarnitine | C3 | Arginine | Arg |
| Malonylcarnitine | C3DC | Asparagine | Asn |
| Butyrylcarnitine | C4 | Aspartate | Asp |
| Methylmalonylcarnitine | C4DC | Citrulline | Cit |
| Hydroxybutyrylcarnitine | C4OH | Cystathionine | Cystath. |
| Isovalerylcarnitine | C5 | Cystine | Cystine |
| Tigloylcarnitine | C5:1 | Glutamate | Glu |
| Glutarylcarnitine | C5DC | Glutamine | Gln |
| Hydroxyisovalerylcarnitine | C5OH | Glycine | Gly |
| Hexanoylcarnitine | C6 | Histidine | His |
| Adipoylcarnitine | C6DC | Isoleucine | Ile |
| Octanoylcarnitine | C8 | Leucine | Leu |
| Octenoylcarnitine | C8:1 | Lysine | Lys |
| Hydroxyoctanoylcarnitine | C8OH | Methionine | Met |
| Decanoylcarnitine | C10 | Ornithine | Orn |
| Decenoylcarnitine | C10:1 | Phenylalanine | Phe |
| Decadienoylcarnitine | C10:2 | Proline | Pro |
| Hydroxydecanoylcarnitine | C10OH | Hydroxyproline | ProOH |
| Lauroylcarnitine | C12 | Serine | Ser |
| Lauroleoylcarnitine | C12:1 | Taurine | Tau |
| Myristoylcarnitine | C14 | Threonine | Thr |
| Tetradecenoylcarnitine | C14:1 | Tryptophan | Trp |
| Tetradecadienoylcarnitine | C14:2 | Tyrosine | Tyr |
| Hydroxymyristoylcarnitine | C14OH | Valine | Val |
| Palmitoylcarnitine | C16 | | |
| Palmitoleoylcarnitine | C16:1 | | |
| Hexadecadienoylcarnitine | C16:2 | | |
| Stearoylcarnitine | C18 | | |
| Oleoylcarnitine | C18:1 | | |
| Linoleoylcarnitine | C18:2 | | |
| Linolenoylcarnitine | C18:3 | **Important note** Taurine amino acid is abbreviated with Tau. This is not related to Tau-protein. | |
| Hydroxystearoylcarnitine | C18OH | | |
| Eicosaenoylcarnitine | C20:1 | | |
| Eicosadienoylcarnitine | C20:2 | | |
| Eicosatrienoylcarnitine | C20:3 | | |
| Arachidonylcarnitine | C20:4 | | |

### Text S8: Untargeted metabolomics data processing from plasma and blood samples (prepared from the acylcarnitine method).

The plasma and blood samples were prepared according to the acylcarnitine method (see supplementary text S5). The analysis included five technical replicates for both preparations (liquid and filter paper extraction). Raw data was uploaded in Metaboscape 2021b (Bruker Daltonics, Bremen, Germany). Processing was done in positive mode with the timsTOF T-Rex 3D algorithm including the following parameters:

Min. # of features for extraction = 0; Min. # of features for result = 0; Feature type = intensity; Intensity threshold = 500; Scan range = 20 – 1300 m/z; RT range = 0 – 12 min; Mass recalibration = activated with auto-detection mode (sodium formiate positive).

Ion deconvolution: polarity = positive; EIC correlation = 0.8; Primary ion = [M+H]+; Seed ions = [M+Na]+; Common ions = [M+H-H2O]+

Annotations: obtained by using an analyte list (inhouse), MoNA-LipidBlast and Massbank-NIST-June2023 libraries with the following parameters: m/z = 0.5 (narrow) to 1.0 (wide); RT = 0.25 (narrow) to 0.4 (wide); mSigma = 50 (narrow) to 1000 (wide); MS/MS score = 900 (narrow) to 400 (wide). The RT scoring was only applied for the analyte list while the MS/MS score was only applied for the databases.

In plasma, 21742 features and 202 annotated compounds (with internal standards) were found, while in whole blood 16777 features and 204 compounds were identified. From Metaboscape, only the annotated compounds were exported. For the PCA analysis, the data was imported to GraphPad Prism and then a PCA calucation was performed. Standardizing (mean of 0 and SD of 1) was activated and principal components (PCs) selected on percent of their total explained variance (together: 75% explained). For the volcano plot, respective data files were uploaded to the Metaboanalyst 6.0 online platform. No feature filtering was applied, however for normalization of intensity responses the auto-scaling was performed. A volcano plot analysis was used to generate the fold change (FC), log2(FC), the significance p-value and –log10(p-value) parameters. This data was extracted into GraphPad Prism to generate volcano plots. Features with a FC of < 0.5 (blue) or > 1.5 (red) are coloured (Fig. 3, main article).

### Text S9: Clinical DBS metabolomics study (229 samples)

Sample preparation and LC-MS measurement

All samples were prepared by punching out a DBS spot of 6.00 mm diameter and placing them into respective 1.5 mL screw-cap vials. Upon preparation, they were stored at – 20 °C. The samples were prepared by using the amino acid extraction scheme (given under Supplementary Text S6), however the extraction solvent was changed to [MeOH:ACN:$H_2O$] (40:40:20 v/v/v) and the total end volume changed to 300 µL (200 µL solvent addition after ISTD). A pooled QC was prepared by mixing an aliquot of 30 µL from all samples together. Samples were measured within a single batch. The pooled QC was injected after every ten samples and blanks were included for carryover assessment. The measurement was performed in HILIC mode (amino acid settings, described in chromatographic conditions) with an injection volume of 5 µL, however an extended scan range of 20 – 1300 m/z and slightly altered MS parameters were utilized (Table S2).

LC-MS data processing

The raw data processing was performed in Metaboscape 2021b (Bruker Daltonics, Bremen, Germany). The batch was imported and processed with the T-Rex 3D LC-QTOF algorithm in positive mode with the following parameters:

- Min. # of features for extraction = 25; Presence of features in minimum # of analyses = 25; Intensity threshold = 300; Min. # of peaks = 6; Recursive feature extraction = enabled (4 spectra); Scan range = 20 – 1300 m/z, RT range = 0.5 – 10 min.
- Ion deconvolution: EIC correlation = 0.95; Primary ion = $[M+H]^+$; Common ions = $[M+H-H2O]^+$; Seed ions = $[M+Na]^+$
- Recalibration: activated; Detection of calibration segment = auto

With this processing, a final list comprising of 48350 buckets (= features) was obtained. For the annotation of compounds, an analyte list was used together with two databases (MoNA-export-LipidBlast; Massbank-NIST-June-2023). For the analyte list, the scoring parameters were as followed: m/z narrow = 0.5 mDa, m/z wide = 1.0 mDa; RT narrow = 0.25 min, RT wide = 0.4 min; mSigma narrow = 100, mSigma wide = 1000. For the databases, the scoring parameters were as described above, however the RT scoring was replaced by the MS2 scoring: MS/MS narrow = 850, MS/MS wide = 400.

As a next step, the common MS background contamination ion list from Fisher Scientific[1] was used to eliminate contaminations, including various phthalate-, polyethylene glycol- and tributylphosphate compounds.

With this approach, 223 annotations were generated (239 including internal standards). A list comprising of these 223 annotations with all samples was exported and further processed for blank carryover-assessment and QC signal drift correction.

The annotation quality is separated into two levels, with level 1 showing a high security based on RT matching with (isotopic) standards and/or a high MS2 spectra fit. Level 2 annotations have a lowered security and are based on MS1 accuracy and/or implementation from a published HILIC metabolomics analyte list with a good relative RT fit to arginine between both methods.

Post-processing data corrections

As a first post processing step, in order to filter out features with high carryover, the signal intensity from a blank measured within the sequence was used and compared against the mean signal from all samples before. For the majority of analytes, the carryover was 0%, and features with a carryover of > 15.0% were deleted, which was true for 25 features. The two features of Arginine-ISTD were also combined to one bucket, yielding 214 features. Some other features were deleted: Dopamine (no RT, MS2 match), LPC 18:2 (double), Hydroxyisovalerylcarnitine (double eluting peak). This yields 195 final annotations.

In order to reduce a signal intensity drift which was identified with the pooled QC samples, a SERRF[2] algorithm (systematical error removal using random forrest) was utilized using the online tool. The result of the SERRF approach is shown on the next page by five representative metabolites indicating how the pooled QC raw signal drift was corrected to an almost perfect linear behaviour (compare Fig. S1-S2).

[1] https://beta-static.fishersci.ca/content/dam/fishersci/en_US/documents/programs/scientific/brochures-and-catalogs/posters/fisher-chemical-poster.pdf
[2] https://slfan2013.github.io/SERRF-online/

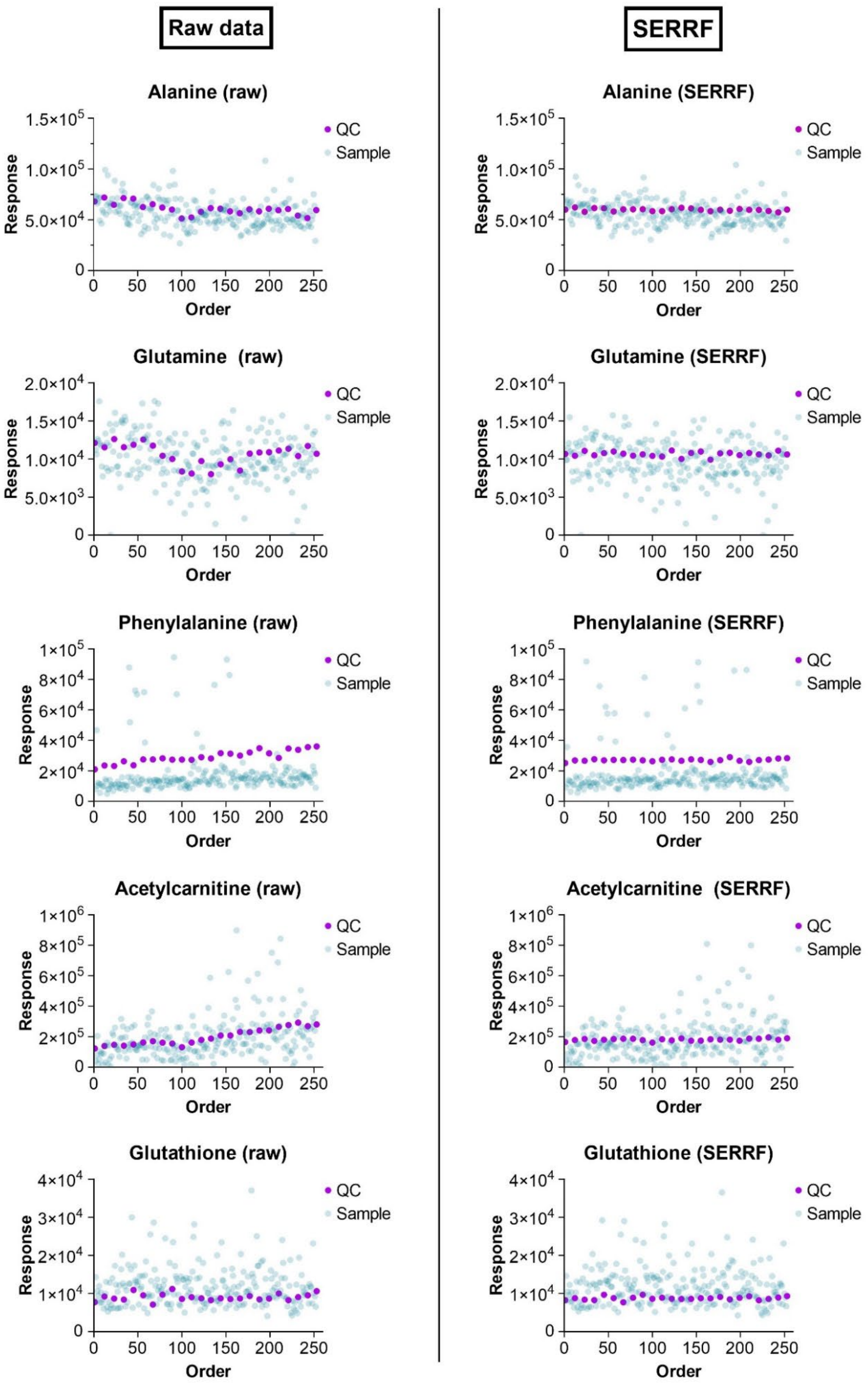


**Figure S1:** The SERRF machine-learning algorithm corrects various signal intensity drifts excellently, which occurred over the measurement.

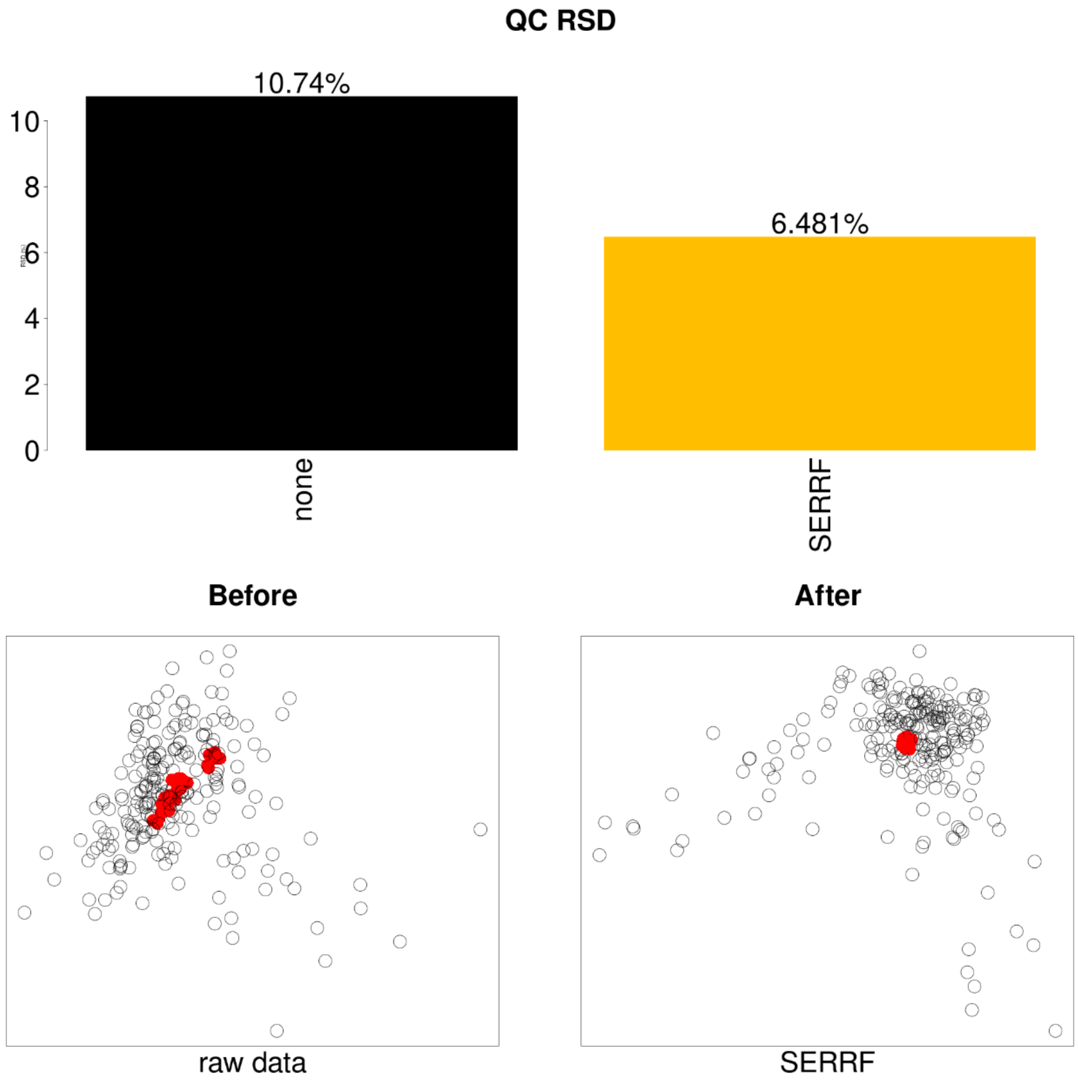


**Figure S2:** Overall performance of SERRF QC batch correction. The mean QC RSD was reduced to 6.5% (originally: 10.7%) and the QCs clustered in a single spot (bottom, right).

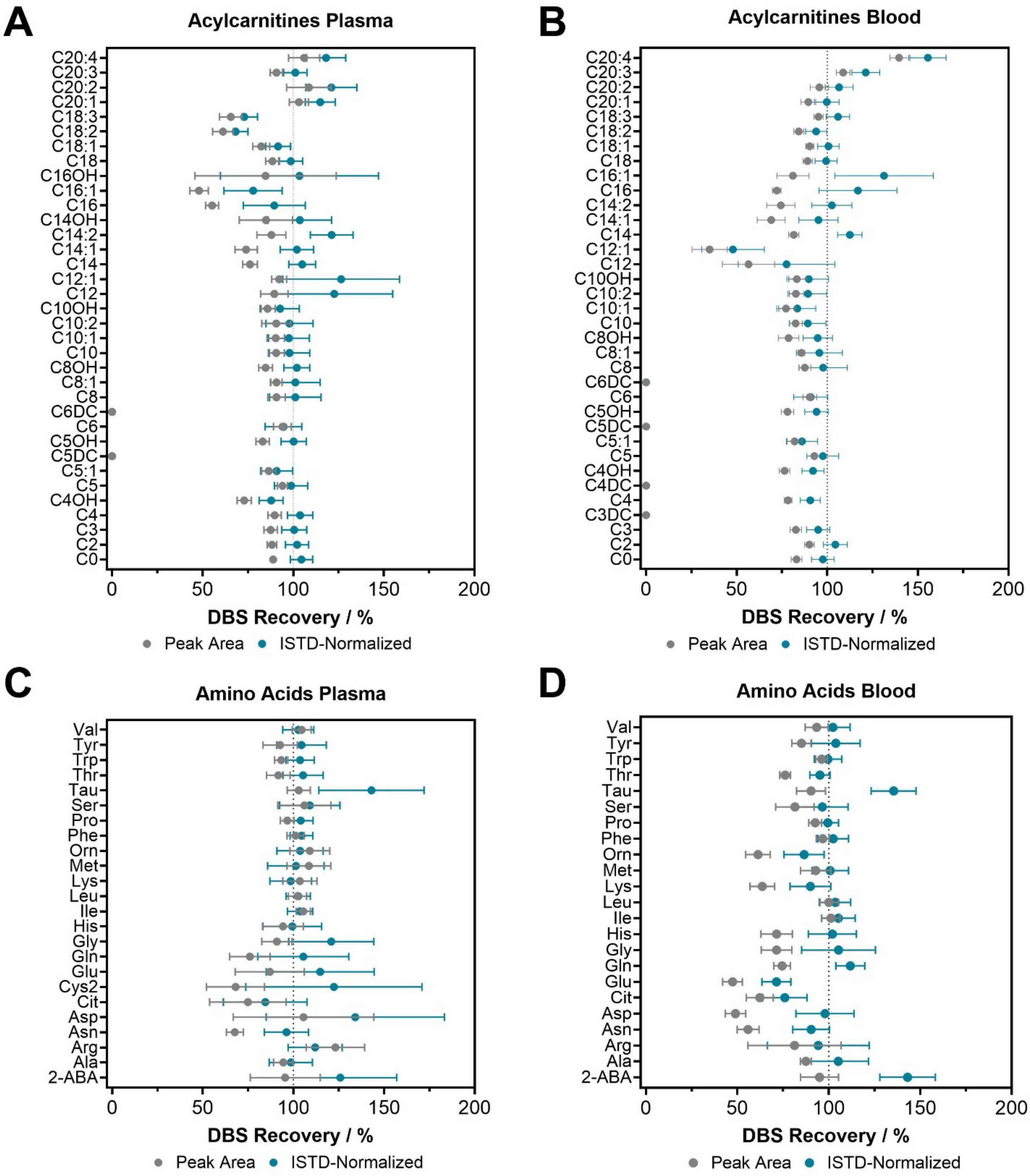


**Figure S3:** Vertical point plots depict the mean ± SD peak area and ISTD-normalized recovery for acylcarnitines in plasma (A) and whole blood (B). Amino acid results are given for plasma (C) and whole blood (D).

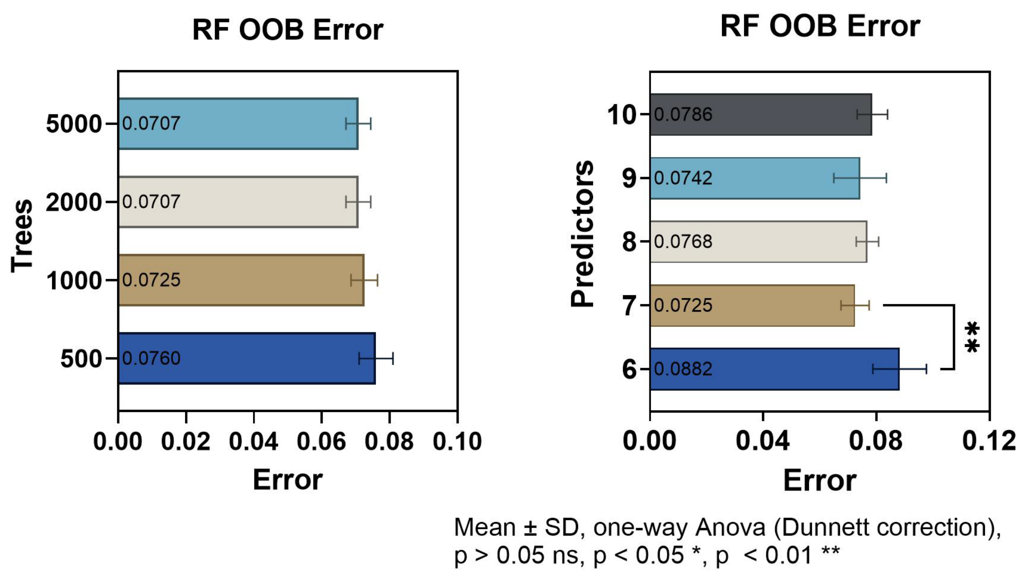


**Figure S4:** Influence of the number of trees and predictors in a random-forest model towards the overall out-of-bag (OOB) error.

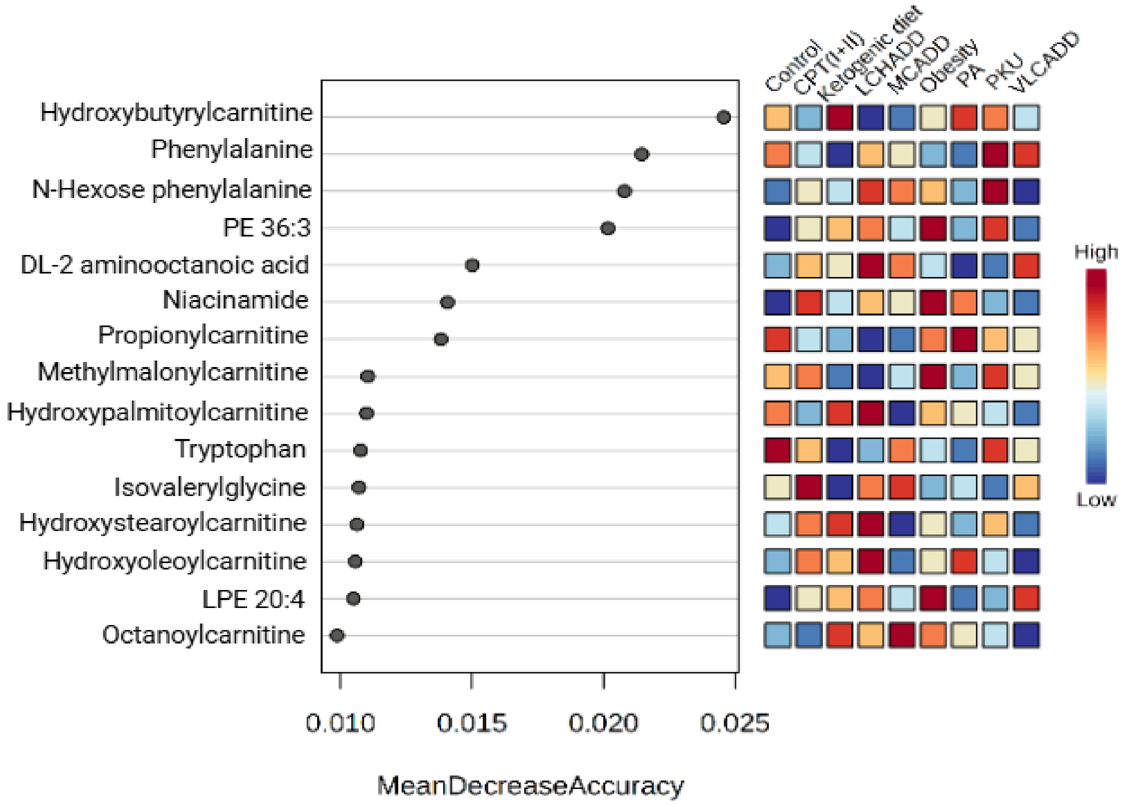


**Figure S5:** Feature ranking plot of a Random-Forest (RF) calculation further depicting the importance of features within specific clinical indications (right). This calculation is based on the standard dataset (no added diagnostic ratios). Due to a certain randomness of RF, this calculation shows features in minimal different ranks than shown in Fig. 7E.
